# Origin and phylogenetic status of the local Ashanti Dwarf pig (ADP) of Ghana based on evidence from mtDNA analysis, MC1R, and Y-chromosome haplotypes

**DOI:** 10.1101/066209

**Authors:** R. Osei-Amponsah, B.M. Skinner, O.D Adjei, J Bauer, G Larson, N.A Affara, C.A Sargent

## Abstract

The Ashanti Dwarf Pig (ADP) of Ghana is an endangered pig breed with hardy and disease resistant traits. Characterisation of animal genetic resources provides relevant data for their conservation and sustainable use for food security and economic development. We investigated the origin and phylogenetic status of the local ADP of Ghana and their crosses with modern commercial breeds based on mtDNA, *MC1R* and Y-chromosome sequence polymorphisms, as well as genome-wide SNP genotyping.

The study involved 164 local pigs sampled from the three agro-ecological zones of Ghana. Analyses of the mitochondrial D-loop region and Y-chromosome sequences revealed both European and Asian genetic signatures, with differences between the geographical zones. Black coat colour is the most predominant within the breed, with dominant black *MC1R* alleles of both Asian and European origin contributing. European alleles for spotting are present at a low frequency in the sample set, and may account for the occurrence of spotted piglets in some APD litters. PCA analysis of SNP data revealed a strong location and breed effect on clustering of local Ghanaian pigs. On a global level, Ghanaian local pigs cluster closely with European pigs of commercial origin, such as the Large White.

The presence of both European and Asian contributions, with differences between geographical zones probably reflects trading and colonial influences.. Understanding the effects of admixture on important adaptive and economic traits of the ADP and other local breeds in Africa is critical for developing sustainable conservation programmes to prevent the decline of these genetic resources.

## Introduction

Pigs (*Sus scrofa*) display enormous phenotypic diversity in terms of shape, colour, size, production and reproduction abilities. Indigenous pig breeds in China, for instance, are well-known for their unique reproductive and lactation performance, good meat quality, strong adaptability and disease resistance traits (Wang *et al.*, 2010).

The Ashanti Black Forest Dwarf Pig of Ghana, commonly called the Ashanti Dwarf Pig (ADP), is a local breed with relatively good adaptive traits raised at the subsistence level in mixed farming systems in Ghana (Tachie-Menson, 1990; Ahunu *et al.*, 1995). The ADP is hardy, tolerant to endemic diseases, survives under poor management conditions and heat stress, and is also able to handle fibrous feeds much better than exotic breeds (Darko and Boadu, 1998; Barnes and Fleischer, 1998). Ghana’s report on Animal Genetic Resources (AnGR) indicates that, apart from the ADP, there are various locally adapted exotic breeds as well as crosses between the exotics and the local ADP (APD, 2003). Thus, in spite of its relatively low cost of production, the ADP has over the years faced a threat from exotic breeds such as the Large White, Landrace and Duroc (Barnes and Fleischer, 1998).

Genetic characterisation of the ADP, including an understanding of its historical origin, remains unknown whilst phenotypic distinctions between purebred and F_1_ crossbred pigs is often not possible because of the dominant black colour of the ADP (Osei-Amponsah *et al.*, 2015). An assessment of the origin and genetic diversity of the ADP and its relationship with exotic breeds will be a major step toward the development of sustainable conservation and improvement programmes to prevent its decline and extinction (Amills, 2011). This will also provide more information on the breed and help in identifying it as a breed critical for preservation (Eggén, 2012).

A number of approaches have been developed to study the origin, genetic variation and unique attributes of animal genetic resources providing valuable information for their conservation.. Mitochondrial DNA (mtDNA) has been used to produce phylogenetic trees at several taxanomic levels, from within species to among orders (e.g., Larson *et al.*, 2005; Haile *et al.*, 2010). MtDNA is maternally inherited, haploid, non-recombining and its evolutionary rate of base substitution is much faster than that of nuclear DNA (Avise, 2000). Thus it can be used to follow the maternal contributions within the porcine domestication process (Kim *et al.*, 2001: Fang *et al.*, 2006). In contrast, fragments of sequence from the Y chromosome have been analysed to study paternal lineages in domesticated pigs (Ramos *et al.*, 2009; Cliffe *et al.*, 2010). These studies show that there is more than one porcine Y chromosome lineage in domestic breeds and European wild boar based on the combinations of major alleles on the non-recombining portion of the Y chromosome (NRY). Four different major Y chromosome populations can be identified based on the *SRY* gene which is responsible for sex determination in mammals (Gubbay *et al.*, 1990; Sinclair *et al.*, 1990; Koopman *et al.*, 1991). Mitochondrial, *SRY* and additional NRY sequence polymorphisms are described in this study as part of the analysis of the genetic origins of the ADP.

Coat colour is another important trait in characterization of animal breeds (Switonski *et al.*, 2013) and among the genes affecting this trait, the melanocortin receptor 1 (*MC1R*) locus is the most consistently polymorphic (Margeta *et al.*, 2013). *MC1R* is primarily expressed in melanocytes and plays a key role in melanogenesis by determining the switch between production of red/yellow pheomelanin and dark eumelanin (Robins *et al.*, 1993). Mutations in the *MC1R* gene affect coat colour in pigs (Kijas *et al.*, 1998; 2001); loss-of-function mutations are associated with recessive red coat colour or spotting, whereas dominant black colouring is linked with mutations affecting *MC1R* signalling (Margeta *et al.*, 2013). As black is the dominant colour of the ADP, sequences from ADP or local cross-bred animals, were compared against the porcine reference genome and matched to haplotypes defined in Fang *et al.* (2009). The distribution of Asian and European dominant black alleles differs by region and by classification (ADP or cross-breed), with both European and Asian haplotypes represented.

Based on the selected DNA regions, results suggest the ADPs have both European and Asian ancestry, with a difference between animals from the north and south of the country. A further study performed using the Illumina Porcine SNP60k BeadChip provided genome wide data for comparison with the deposited genotypes of animals collected across the globe (Burgos-Paz *et al.*, 2012). Using a PCA analysis, local pigs appear to lie on the edge of the cluster of commercial European pigs such as Large White and Landrace. An F_ST_ analysis identified regions of the genome differing between ADPs and European commercial pigs, Chinese pigs and Duroc. Within these regions, we identified intervals with the genes *LCORL* and *PLAG1*, previously associated with height and body size in other species (reviewed by Takasuga, 2016). This comparison provides targets for future sequencing approaches.

## Results

For investigation of the genetic relationships, and evaluation of local Ghanaian pigs, 165 animals were sampled from the agro-ecological zones as shown in figure 1. A full list of samples can be found in Supplementary_Table_S1.xlsx.

**Figure 1:**
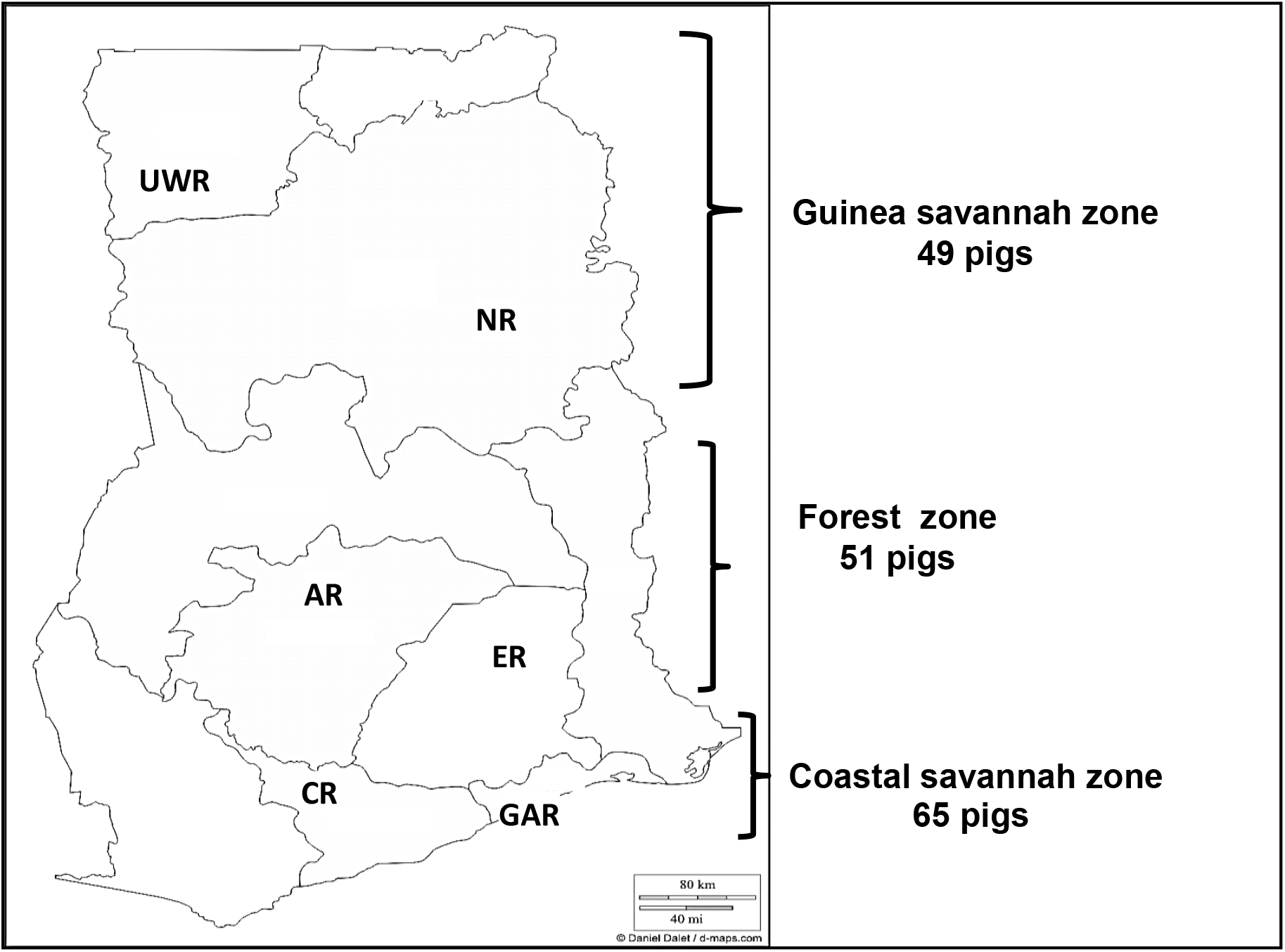
Map of Ghana showing regions where pigs were sampled. (UWR = Upper West Region; NR = Northern Region; AR = Ashanti Region; ER = Eastern Region; CR = Central Region; GAR = Greater Accra Region)

### Mitochondrial Haplotype Analysis of Sequences

Mitochondrial DNA sequence analyses of 140 animals were used to develop a Bayesian phylogenetic tree (Figure 2). Ghanaian local pigs clustered into 2 clades made up of 14 haplotypes, of which 8 clustered with European and 6 with Asian *Sus scrofa* haplotypes. Six of the sequences had perfect matches to existing entries in the GenBank database, whilst the remaining 8 were unique to this study (see Supplementary_Table_S2.xlsx). The majority of sequences from animals sampled in this study fell into three haplotypes (haplotypes 3, 8, 13: 75%). Two fall into the European clade (haplotypes 3, 13) and one into the Asian clade (haplotype 8).

**Figure 2:**
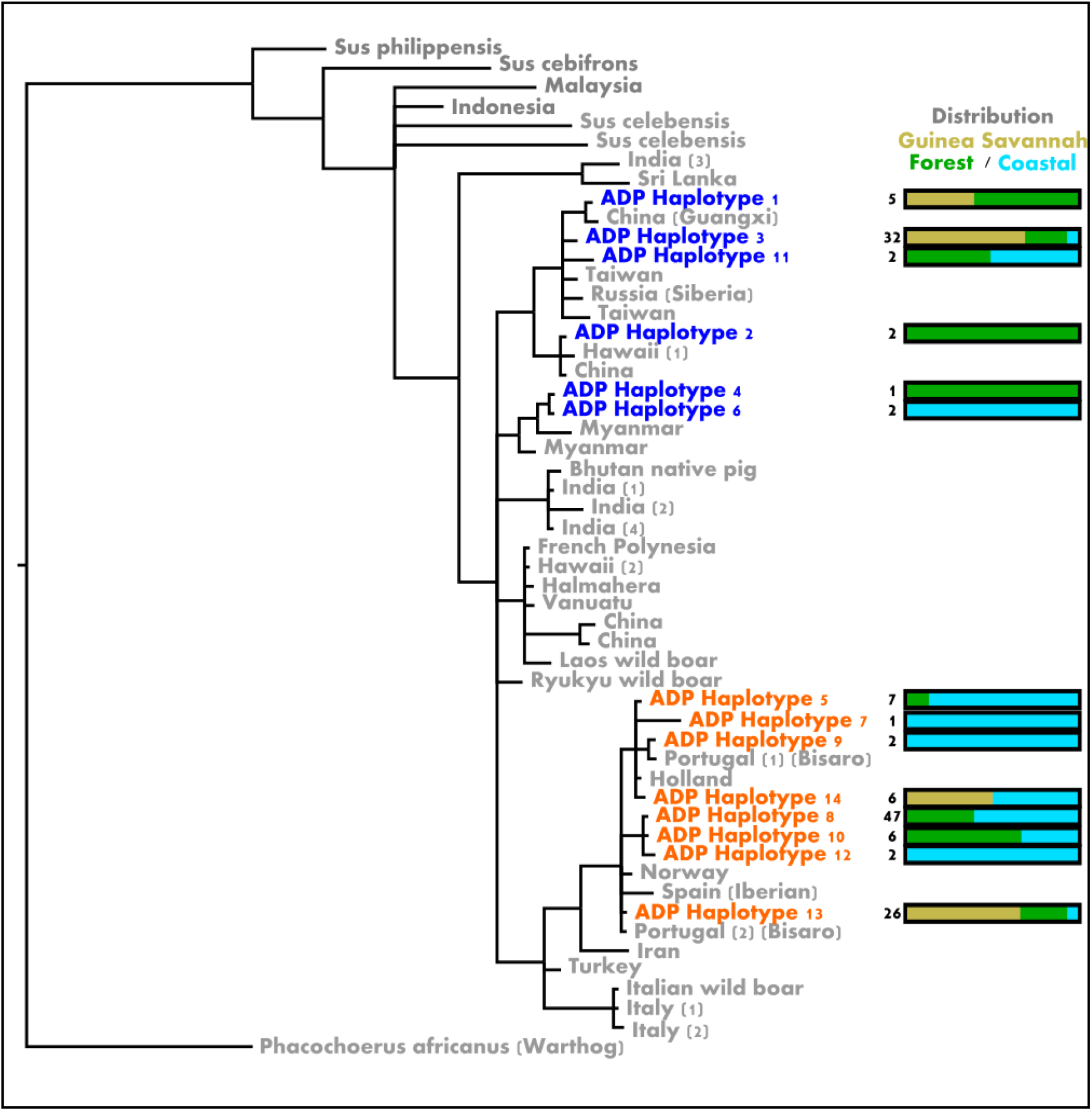
A Bayesian phylogenetic tree based on analysis of the mitochondrial D-loop region. The African warthog was used as an outgroup. The panel shows the local haplotypes (in blue) clustering with sequences of Asian origin, and those (in orange) clustering with the sequences of European origin. To the right of each haplotype, the number beside the bar gives the total for individuals that share the haplotype, and the bar shows the regional distribution.

#### Mitochondrial genetic distances between ADPs and other pig breeds

The clustering of haplotypes into European and Asian clades was confirmed using base substitution data, with smaller genetic distances observed between the two major European haplotypes identified in this study compared to the genetic distances between the major Asian and European haplotypes (Table 1). Globally, the mitochondrial sequences from the ADP of Ghana were closer to European than Asian sequences, as shown in Table 2.

**Table 1:** Mean genetic distances between the three predominant mtDNA haplotypes representing the majority of the animals sampled (see figure 3). The number of base substitutions per site from averaging over sequence pairs between haplotype groups are shown. Standard error estimate(s) are shown in the upper diagonal (italicised). Haplotype 3 is within the Asian clade; haplotypes 8 and 13 are within the European clade.

| ADP haplotype | 3 | 8 | 13 |
| --- | --- | --- | --- |
| 3 |  | *0.008* | *0.008* |
| 8 | 0.026 |  | *0.003* |
| 13 | 0.026 | 0.004 |  |

**Table 2:** Within breed genetic distance of various *Sus* groups from mtDNA comparisons. The number of base substitutions per site from averaging over all sequence pairs between groups is shown. Standard error estimate(s) are shown in italics in the upper diagonal; ADP = Ashanti Dwarf pig.

| Breed | ADP | Asian | European | Pacific | *Sus* outgroups | African Warthog |
| --- | --- | --- | --- | --- | --- | --- |
| ADP |  | *0.005* | *0.003* | *0.005* | *0.007* | *0.013* |
| Asian | 0.027 |  | *0.005* | *0.004* | *0.006* | *0.014* |
| European | 0.018 | 0.027 |  | *0.005* | *0.007* | *0.014* |
| Pacific | 0.026 | 0.022 | 0.025 |  | *0.006* | *0.014* |
| *Sus* outgroups | 0.043 | 0.038 | 0.042 | 0.033 |  | *0.012* |
| African Warthog | 0.095 | 0.099 | 0.096 | 0.095 | 0.090 |  |

### SRY Sequencing and chromosome Y haplotypes

*SRY* sequences were obtained from 33 males: 21 from animals selected from the designated ADP populations and 12 from local crossbred animals. All but two animals had the *SRY* haplotype found to predominate in European animals as described in Cliffe *et al.* (2010). Two animals, one from the ADP population, and one from the crossbred population, both of the Ashanti region (AR), had haplotypes previously observed in Tamworth and some Asian breeds (Cliffe *et al.*, 2010). No novel *SRY* sequences were detected.

Thirty three males (22 ADP, 2 exotic and 9 local crossbred) were identified from the genome wide genotyping panel. One individual had previously been classified as female (animal 163), but appears to be karyotypically male based on both chromosome X and chromosome Y SNP data. One individual originally assigned as male in the records appeared to be female, based on the SNP data (animal 149). All other animals were genotyped in agreement with the recorded genders. Of the genotyped panel, 22 DNA samples from males were in common with those selected for *SRY* sequencing. The extended Y chromosome haplotypes were in agreement with the previously identified *SRY* haplotypes for all of these animals (Supplementary_Table_S3.xlsx).

### Coat Colour in Ghanaian local pigs

The ADP of Ghana, although predominantly black, displays other coat colours and patterns. These can be observed even in the offspring of selected “purebred” black-coated ADPs, suggesting the interplay of multiple coat colour genes in the phenotypic outcome (Supplementary_Figure_S3.pdf). Analysis of the collected data shows that the local animals include spotted, white or belted animals, as well as those with black coats (Supplementary_Figure_S4.pdf).

Since black coat colour is largely determined in mammals by the dominant alleles of the *MC1R* gene, the nature and origins of these alleles in Ghana were investigated through DNA sequence analysis.

### MC1R gene distribution in Ghanaian local pigs

In total, 86 Ghanaian pigs were fully or partially sequenced for the *MC1R* gene and promoter region. Control sequences from purebred red Duroc, and Large White animals were included and compared with the reference porcine genome.

The identifiable haplotypes for *MC1R* represented the European 301 (E^D2^; dominant black), Asian 201 (E^D1^; dominant black), European 501 (E^P^; spotting) and European 401 (red) variants of the gene. Sequence signatures for the respective promoters were confirmed by BLAST analysis of our data with *MC1R* entries in the public databases. Using these four major identifiable haplotypes, genotypes were predicted in the Ghanaian population, (Supplementary_Table_S4.xlsx). For animals with incomplete or inconclusive sequence information, alleles were assigned of ‘Asian’ or ‘European’ origin based on the available data. Four animals (5, 143, 150, 158) have single base sequence deviations that do not fit the previously reported alleles. These may represent local variants of the major haplotypes, especially as the same sequence data are observed twice in animals from the Upper West Region (UWR; 150 and 158): further work with additional samples from local Ghanaian pigs is required to confirm these observations.

As with the mitochondrial data, the occurrence of Asian alleles was higher in animals from the Guinea Savannah zone than those from the other regions. Further discussion of the distribution of alleles is given in Supplementary Information.

### Genotyping Analysis

Genotyping data were used to perform comparisons using PCA analyses, genome wide levels of homozygosity, and an F_ST_ comparison of the ADPs against other world pig breeds.

The PCA analysis of local Ghanaian animals is shown in Supplementary_ Figure_6A.pdf. ADPs form two main groups, based on geographical location: those from the Guinea Savannah zone are the most tightly grouped, followed by those from the Coastal zone (particularly GAR), with the Forest zone animals being more diffusely scatted across the plot. In general, the crossbred pigs tended to be closer to the exotic (European breed) animals, as might be predicted based on known introduction of European breed genetics. When data from other world pig breeds are included, the Ghanaian pigs cluster more closely with European commercial breeds than with Asian pig breeds (Supplementary_ Figure_6B.pdf).

#### F_ST_ analysis

We performed an F_ST_ analysis between two ADP subgroups and between the ADPs and European and Chinese breeds. The first ADP subgroup was selected from the Guinea Savannah zone, and formed a distinct cluster in PCA analysis (Supplementary_ Figure_6A.pdf set 1).

The second subgroup came from the Coastal zone (Supplementary_ Figure_6A.pdf set 2). The two major subgroups of ADPs (Guinea Savannah and Coastal) comprise 17 animals each.

Genome averaged F_st_ values are consistent with the mitochondrial data: the ADPs are genetically more similar to European Commercial breeds than to either Duroc or Chinese breeds (Table 3)

**Table 3:** Genome average F_ST_ values for each of the two ADP populations (GSZ: Guinea Savannah zone; CZ, coastal zone) tested against European commercial breeds, Duroc and Chinese breeds.

|  | set 1<br>(GSZ) | set 2<br>(CZ) |
| --- | --- | --- |
| subset 2 | 0.131 |  |
| European | 0.085 | 0.105 |
| Chinese | 0.334 | 0.319 |
| Duroc | 0.196 | 0.191 |

Comparison of ADPs against European commercial breeds may help to identify regions that distinguish these local pigs from their most similar relatives. Forty two genomic intervals contain SNPs with high F_ST_ (SNP F_ST_ in 99^th^ percentile, within a 100kb window with average F_ST_ greater than the 95^th^ percentile) between both ADP subgroups compared independently against the European commercial breeds. The intervals and gene content are summarised in Supplementary_Table_S5.xlsx, which also indicates regions of difference against Chinese, Duroc, and between ADP groups. Notably, several of the genes in the regions distinguishing ADPs from European breeds relate to skeletal morphology and body size (e.g. *LCORL, IMPAD1, STIM2*; for *LCORL* see Supplementary_Figure S5.pdf), and two regions have genes related to melanin production (*KIT, TRPM1*).

Few significant categories of genes clusters were identified using the DAVID analysis, although genes identified at the peak SNP of F_ST_ regions in both ADP sets against other pig populations can be grouped according to human gene-disease associations (Supplementary_Tables_S6 & S7.xlsx). Here, cardiovascular and haematological functions are the most represented association classes with genes from the ADP comparison against the European breeds. Comparisons with the Duroc, and between the two ADP groups, identify genes that relate to behaviours (e.g. chemical dependency) in humans, and notably the same genes are also implicated in metabolic functions. No groupings were identified in the peak gene list from ADP comparisons with Chinese breeds.

## Discussion

The Ashanti Dwarf Pig (ADP) has often been held as an example of a local pig breed with important genetic characteristics worthy of preservation and conservation. In this study, we show that many of the animals described by local Ghanaian farmers as ADP in fact derive from pigs of different ancestries, and the pure Ashanti pigs themselves may exist in far fewer numbers than previously thought.

Although the history of local African pig breeds is largely unknown and controversial (Blench, 1999), they have been reported to display both European and Far Eastern genetic signatures with a west versus east geographical distribution (Ramirez *et al.*, 2009). Africa was explored and colonized by the Europeans in the 15^th^ century onwards which may explain why many pigs demonstrate genetic signatures mostly of European origin (Ramirez et al., 2009). The presence of Asian signatures could be attributed to the ancient introduction of Indian and Far Eastern livestock to the Eastern coast of Africa (Hamite *et al.*, 2002; Muchadeyi *et al.*, 2008). However, Europeans imported a few Asian breeds during the 18^th^ and 19^th^ centuries and modern European breeds are on average 30% Asian genetics (White *et al.*, 2011; Bosse *et al.*, 2014). Thus more recent introductions may also account for the presence of Asian haplotypes in local Ghanaian pigs.

### Mitochondrial sequences

Analysis of SNP polymorphisms in the D-loop sequence of mtDNA is widely used to study genetic and phylogenetic relationships within population of farm animals (Molnàr *et al.*, 2012). MtDNA has been used to describe variation in putative wild ancestor and modern domestic livestock populations including pigs (Ursing and Arnason, 1998; Bruford *et al.*, 2003; McCann *et al.*, 2014). In other studies, significant differentiation between European and Chinese domestic pigs has also been revealed by mtDNA analyses (Giuffra *et al.*, 2000; Okumura *et al.*, 2001, Watanobe *et al.*, 2001, Kim *et al.*, 2002). Therefore, in order to get an overview of the phylogenetic information content, our sequences were aligned and compared with sequences from other European and Asian swine breeds. The presence of both European and Asian MtDNA haplotypes in all six regions sampled indicates an admixed local pig population in Ghana. However, there is a distinct gradient with Asian sequences more common in the Guinea Savannah Zone, and European sequences on the coast. Here, the predominance of European mtDNA haplotypes may be explained by the influence of Europe on Ghana as a result of colonization and imports of European breeds by the Ghanaian government in the recent past to boost pig production.

### Y chromosome Analysis

Results of the present study on the paternal ancestry of local pigs in Ghana show that almost all the tested samples had Y chromosome signatures commonly found in multiple European breeds, with only two animals having *SRY* sequence data comparable to the Tamworth. The extended Y haplotype signature observed in Tamworth was confirmed in the single male genotyped on the Illumina panel. However, such Y chromosome data cannot rule out non-European origins or admixture, since the observed Y lineages are also found in Asian pig breeds (Cliffe *et al.*, 2010).

### Coat colour

Coat colours of domestic animals mainly represent human needs or cultural preferences and result from strong artificial selection and domestication bottlenecks (Innan and Kim, 2004). A series of alleles of the *MC1R* gene have been identified in pigs. Kijas *et al.*, (1998, 2001) reported that six major allelic variants corresponded to five different patterns of expression. The wild-type (*E*^+^) allele, found in wild boars, corresponds to haplotypes 0101 (*MC1R*\*1) and 0102 (*MC1R*\*5). The dominant black colour results from two different mutations, each of which evolved independently in Asia and in Europe. Large Black and Black Meishan pigs carry the Asian dominant black allele (*E^D1^*) corresponding to 0201 (*MC1R*\*2: *Leu102Pro, Val95Met*); whereas Hampshire pigs carry the European dominant black allele (*E^D2^*) corresponding to 0301 (*MC1R*\*3: *Asp124Asn*). Landrace, Yorkshire and Pietran carry the spotting allele (*E^P^*) corresponding to 0501 (*MC1R*\*6: *nt67insCC, Asp124Asn*) which causes black spotting on a red or white background (Fang *et al.*, 2009; Margeta *et al.*, 2013). The black spots are attributed to the recovery of *MC1R* protein function via somatic mutations leading to restoration of the open reading frame (Kijas *et al.*, 2001). The Duroc breed carries the red coat colour allele (e) corresponding to 0401 (*MC1R*\*4: *Ala164Val* and *Ala243Thr*) (Dun *et al., 2007*).

Unlike Chinese domestic pigs which displayed low diversity of *MC1R* (Yang *et al.*, 2010), Ghanaian local pigs show considerable variation in this gene. This indicates either acquired mutations (Kijas *et al.*, 1998) or an admixed population. Interestingly, the distribution of Asian and European dominant black variants broadly mirrors that of the mitochondrial data: ADP samples from the Guinea Savannah show a higher percentage of the Asian allele (almost 70% of all *MC1R* variants), and ADP samples from the coast a higher percentage of the European allele (almost 40% of all variants). The *E^P^* allele, is also common in the local Ghanaian population, mainly in crossbred animals, but also in those classified as ADPs. In general, local crossbred pigs carrying at least one *E^P^* allele had white or spotted coats, whilst crossbred pigs of other genotypes were black, belted and white. The presence of white coated crossbred animals genotyped here as homozygous for dominant black alleles is also indicative of the influence of epistasis between coat colour loci, for example the dominant white *KIT* allele, in the local population.

### Genome-wide analysis

The Illumina Porcine SNP60k BeadChip (Ramos *et al.*, 2009) has been used to identify SNP associated with, for example, reproduction traits in the Finnish Landrace pig breed to provide valuable candidates for possible marker-assisted selection (Uimari *et al.*, 2011); detail a genome wide overview of “indigenous” local pig populations (Burgos-Paz *et al.*, 2012) and determine population structure, linkage disequilibrium (LD) pattern and selection signature in Chinese and Western pigs (Ai *et al.*, 2013). In this study, we have used the same chip to investigate the polymorphisms present in the local Ghanaian pig populations. The principal component analysis (PCA) of the local pig populations revealed distinct clustering based on the origin of the samples from across the three agro-economical zones. Based on biological knowledge, PCA1 probably represents geographical distribution, and PCA2 genetic distance. At the whole genome level, ADP from Greater Accra may be genetically similar to animals from the Guinea Savannah zone. Crossbred animals are generally closer to the exotics (European commercial breeds), and ADPs from the forest zone (AR and ER) and the CR seem to be the most diverse. In the PCA analysis that used data from breeds collected from Europe and China, all Ghanaian local pigs, irrespective of classification, overlapped and clustered with the commercial European breeds represented by the inclusion of data from Large White and Landrace samples.

### F_ST_ data shows genes that may underlie the phenotypic distinctiveness of ADPs

The F_ST_ analysis reveals regions of the genome that have greater or lesser genetic similarity between populations. The regions we identified distinguishing ADPs from European commercial breeds contain intervals with genes known to relate to body size and shape, previously reported as genomic intervals undergoing selection in pigs and other mammals (Rubin et al, 2012; Takasuga, 2016). These may help explain the short stature of the ADPs compared to other breeds, and will be useful targets for more detailed study.

We propose that pigs from the southern part of Ghana should be more admixed due to colonisation, and relatively more activity in terms of trade and movement of people and animals to that part of the country. Correspondingly, pigs from these regions show more variation both in mitochondrial DNA and *MC1R* sequences than those from the northern Guinea Savannah. Animals from the Northern part of the country have higher percentages of Asian signature sequences at the selected loci, but it is currently unclear if this is indicative of historic breeding bottlenecks or points to two distinct geographical origins of the animals currently classified as Ashanti Dwarf Black pigs.

## Conclusions

Local ADPs of Ghana display genetic signatures indicative of both European and Asian origins at the loci described here. Although the ADP is nominally a black coated breed, the recent occurrence of spotted piglets in APD litters may be due to epistatic interactions, or a low frequency of the recessive *E^P^* allele in the selected populations introduced through unrecorded crossbreeding with other local pigs. The data presented suggest that morphology alone cannot be used to adequately characterise Ghanaian local pigs. It will be necessary to sample a larger population of local pigs in Ghana to find out how the adaptive and economic traits of the ADP have been affected by crossbreeding, and define allelic variants of value to the longer term animal breeding programme in Ghana.

## Materials and methods

### Samples and DNA extraction

A total of 165 pigs made up of local ADPs, crossbreds and exotic pigs were sampled from six regions in the three agro-ecological zones of Ghana, namely the coastal savannah, forest and guinea savannah zones (Figure 1) between August 2013 and October 2013.

The samples were obtained from a total of 54 local pig farmers and 6 institutional pig farms. Further details are in Supplemental Materials, table S1. The farmers/managers were interviewed to obtain information on the husbandry practices, such as their experience in keeping the local breed, and if they have ever crossbred their local stock with exotics. Ear tissues of sampled pigs were obtained using an ear notcher (Supplementary_Figure_S1.pdf) with assistance from animal production officers, veterinary technicians and extension agents of the Ministry of Food and Agriculture (MOFA). The samples were stored on field in RNAlater (tissue collection stabilization) solution (*Ambion,* USA) and later transported to the Biotechnology Laboratory of the School of Agriculture, University of Ghana, Legon. Genomic DNA was extracted from the ear tissues using the QIAGEN DNeasy Blood and Tissue Kit following the Manufacturers’ protocol after which the quality of the DNA obtained was tested using a spectrophotometer and stored at -80°C. The DNA samples were subsequently transported to the laboratory of the Mammalian Genetics Group of the Department of Pathology, University of Cambridge for sequencing and genotyping.

### DNA amplification and sequencing

#### MtDNA sequencing

A segment of the D-loop region (approximately 680 bp) of the mtDNA was amplified from 140 local animals (81 classified as ADP and 59 classified as cross-bred) using the following primers pairs: L15387 (5'-CTCCGCCATCAGCACCCAAAG-3' forward) and H124n (5'-ATRGCTGAGTCYAAGCATCC-3' reverse) (Larson *et al.*, 2005). PCRs were set up using the manufacturer’s recommended conditions with Qiagen HotStarTaq^®^ DNA polymerase in the presence of buffer Q. All reactions were carried out in 20 μL volumes, with 0.5 μM primers, and 20-50 ng DNA. The touchdown PCR was set at 95°C for 15 min for the initial denaturation and *Taq* activation. This was then followed by 35 cycles at 94°C for 1 min to denature the template DNA, 1 min at the annealing temperature and another 1 min at 72°C The annealing temperature started at 62°C, decreasing by one degree per cycle until 53°C was reached. The remaining cycles were completed at an annealing temperature of 53°C. The samples were held at 72°C for 10 min and chilled at 4°C until removed from the PCR machine.

PCR products were purified following agarose gel electrophoresis using ExoSAP-IT (USB Corporation, USA) following the manufacturer's recommendations Amplicons were sequenced using Big Dye version 3.1 (Applied Biosystems). The sequencing program consisted of 30 cycles of: 96°C for 10 sec, 55°C for 5 sec and 60°C for 4 min. The products were then run on an ABI 3100 capillary sequencer at the sequencing facility in the Department of Biochemistry, University of Cambridge. Traces were edited using Chromas version 2.2 (Technelysium Pty Ltd) before comparing in Sequencer (Genecodes Corporation). Sequences from different animals were also viewed using the MultAlin program (http://prodes.toulouse.inra.fr/MultAlin/MultAlin.html), and within the ClustalW2 program (http://www.ebi.ac.uk/Tools/clustalw2/). Edited consensus sequences and polymorphisms associated with this study are deposited in GenBank under accession numbers KU306949-KU306962.

#### MC1R sequencing

Two primer pairs were used to amplify the majority of the single exon *MC1R* gene. The first pair was MF1 (5' -GTGCGGCGGCTCTGGGCTCCAA forward) and MR1 (5' -CCCCCACTCCCCATGCCTCCTG reverse) whilst the second primer pair was MF1 (5' – GTGCGGCGGCTCTGGGCTCCAA forward) and MR2 (5' – ACACCATGGAGCCGCAGATGAGC reverse). PCRs were carried out in a DNA thermal cycler [Perkin Elmer (Norwalk, CT) 9600] in a total volume of 20 μl containing 25 ng genomic DNA, 1.0 Mm MgCl_2_, 50 Mm KCl, 10 Mm Tris-HCl, pH 8.3, 200 μM dNTPs, 0.5 units AmpliTaq Gold [Perkin Elmer (Norwalk, CT) 9600], and 0.5 μM each of forward and reverse primer. To activate AmpliTaq Gold, initial heat denaturation was carried out at 94°C for 10 min followed by 32 cycles each consisting of 45 sec at 94°C, 45 sec at 53°C and 45 sec at 72°C. The final extension lasted for 7 min at 72°C. Sequencing reactions were purified, run and analysed as above.

#### SRY Sequencing

Primers capturing the entire open reading frame of the *SRY* gene were used to amplify DNA products from male pigs as previously described (Cliffe *et al.* 2010). Samples were purified, run and analysed as above.

### Analysis of mtDNA sequence data

Sequences from the Ghanaian samples were trimmed to remove the amplification primer sequences, then aligned against each other to define 14 unique haplotypes. Reference sequences for each of the haplotypes were taken through the analyses defined below.

Sequence comparisons of the D-loop mtDNA were performed for indigenous Ghanaian pigs and selected porcine mtDNA sequences from the GenBank. The corresponding sequence of the African Warthog (*Phacochoerus aethiopicus*) (GenBank: AB046876) was used as outgroup. Sequences were aligned using Muscle. Evolutionary analyses were conducted in MEGA6 (Tamura *et al.*, 2011). Genetic distances within and between breeds were calculated as the number of base substitutions per site from averaging over all sequence pairs between groups. Standard error estimate(s) were obtained by a bootstrap procedure (500 replicates). Analyses were conducted using the Tamura-Nei model (Tamura and Nei, 1993). The rate variation among sites was modelled with a gamma distribution (shape parameter = 0.33). The analysis involved 53 nucleotide sequences. All ambiguous positions were removed for each sequence pair. There were a total of 573 positions in the final dataset.

A Bayesian phylogenetic tree was constructed from the aligned sequences The analysis was performed in MrBayes, using five million generations with sampling every 5000 generations. Traces were checked in Tracer (http://tree.bio.ed.ac.uk/software/tracer/) and burn-in generation was set at 1000. The final tree was visualised and annotated using FigTree (http://tree.bio.ed.ac.uk/software/figtree/).

### SNP Genotyping and quality control

In this study, genomic DNA of 71 animals were genotyped using the Illumina PorcineSNP60 BeadChip following to the manufacturer’s protocol. One animal was genotypes twice as an internal control. Raw data were visualized and analyzed with the Genome Studio software (Illumina, San Diego, CA, USA).

The SNP genotype calls were exported and loaded in PLINK (Purcell *et al.*, 2007) to perform the PCA analysis. 61565 SNPs were present at the start of the analysis. The filtering parameters were as follows: the maximum missing rate per SNP was set at 10%, minimum allele frequency at 5%, and maximum individual missing rate at 10%. 3305 variants were removed due to missing genotyping data, 6951 due to minor allele frequency threshold, and 4 samples due to individual missing SNP rate. After filtering the 51309 SNPs remained, with 67 individuals successfully typed.

The present raw dataset was also merged with data from a previous study using pig breeds selected from the Americas, Europe and Asia (Burgos-Paz *et al.*, 2013). For this study, based on potential origins of local pigs, historic and current trading routes, a subset of European commercial, Iberian, European wild boar and Asian populations was selected for integration. The two sets were merged resulting in a starting set of 240 individuals with 45673 SNPs in common. The same filters were applied as above, excluding 3 individuals and 1445 SNPs, and leaving 237 individual and 44228 SNPs for the PCA analysis.

Based on the PCA analysis, subsets of tightly grouping ADP populations from the Guinea Savannah and Coastal zones were selected for further characterisation. Each of the two subpopulations comprised 17 animals. Each animal was analysed in PLINK (Purcell *et al.*, 2007) for long stretches of homozygosity, using a sliding window of a minimum run of 100 SNPs. The animals within the same groups were compared for

The male samples were also differentiated by comparing SNPs on their Y-chromosomes to European and Asian boars.

### F_ST_ Analysis

Comparisons of Wright's **F_ST_** s (after Weir and Cockerham,1984) for animals from different geographic regions was performed within PLINK (Purcell *et al.*, 2007) with the 45673 SNPs identified above. The sex chromosome markers, and SNPs which were fixed in any of the populations under comparison were removed. In total, 39848 markers remained for comparison of each of the ADP subgroups against the Chinese and European breeds, and 32793 remained when comparing the two ADP subgroups to each other. The resulting **F_ST_** data was analysed in R. The raw **F_ST_** values were smoothed using a sliding 100kb window, and regions with high **F_ST_** were identified. The threshold to consider a SNP as being of interest were: the individual SNP **F_ST_** was greater than 99% of all F_ST_s, and the smoothed **F_ST_** was greater than 95% of all F_ST_s.

Regions identified by the analysis were scrutinised for gene content in Ensembl. The comparative genomics options allowed identification of any transcripts not fully annotated in the current pig genomic build. Gene lists created from these intervals of interest were further analysed in DAVID v6.8(beta) (https://david-d.ncifcrf.gov/). Genes were classified using known disease classes in humans, to provide the maximum information.

## Data Access

Mitochondrial sequences are deposited under accession numbers KU306949 to KU306962. SNP data is deposited under GEO accession number GSE84604.

## Acknowledgements

The authors are grateful to Cambridge in Africa Research Excellence (CAPREx) for the award of Post-doctoral Fellowship to ROA and the Alborada Trust for funding the Research. The Mammalian Molecular Genetics Laboratory of the Pathology Department, University of Cambridge is acknowledged for hosting ROA during his visit to Cambridge University. We also acknowledge laboratory assistance from Ms. Kerry Harvey, Mrs. Jo Bacon, and Mrs. Kim Lachani of the same laboratory. CAS and NAA are also Fellows of Hughes Hall, Cambridge.

